# Cell-Specific Transcriptomic and Mito-Nuclear Imbalance in Lungs Under Intermittent Hypoxia in Adult Male Mice

**DOI:** 10.1101/2025.08.05.668735

**Authors:** Alexandra Jochmans-Lemoine, François Marcouillier, Marie Martelat, Ynuk Bossé, Dominique K. Boudreau, Sébastien Renaut, Yohan Bossé, Vincent Joseph

**Affiliations:** Centre de Recherche de l’Institut Universitaire de Cardiologie et Pneumologie - Université Laval, Québec, CANADA; Université Laval, Département de Médecine, Québec, CANADA; Université Laval, Département de Médecine Moléculaire, Québec, CANADA; Université Laval, Département de Pédiatrie, Québec, CANADA

## Abstract

Obstructive sleep apnea and its characteristic intermittent hypoxia (IH) are widely recognized as significant contributors to various pulmonary diseases, including asthma, pulmonary arterial hypertension, fibrosis, and chronic obstructive pulmonary disease. While single-cell RNA sequencing (scRNA-seq) has provided valuable insights into cell-type-specific responses to IH, previous studies have primarily focused on post-hypoxic recovery states, leaving immediate molecular responses during active IH exposure unexplored. To address this critical knowledge gap, we investigated real-time transcriptional responses to IH at single-cell resolution in lung tissue using male mice (n=3/group) exposed to either normoxia or IH (30 cycles/h, nadir 6% O_2_, 12 h/day) for 14 days, with tissue collection during active IH exposure. Our analysis revealed pronounced cell-type-specific transcriptional reprogramming, particularly in airway smooth muscle cells (ASMC), arterial endothelial cells (AEC), and lymphatic endothelial cells (LEC). These changes were characterized by enrichment in pathways related to epithelial-to-mesenchymal transition (ASMC, LEC), myogenesis (ASMC), and antioxidant defenses (AEC, LEC). Most cell types demonstrated substantial upregulation of genes encoding mitochondrial complex I-IV proteins and TCA cycle enzymes accompanied by a decreased expression of genes encoded by mitochondrial DNA that was markedly present in LEC, AEC, and cells of the alveolar-capillary unit, revealing a mito-nuclear imbalance. These findings provide novel insights into the immediate cellular responses to IH, showing previously uncharacterized metabolic reorganization that may underlie the development of IH-related pulmonary complications. This improved understanding of early molecular events during active IH exposure advances our knowledge of sleep apnea-related lung pathologies and may inform future therapeutic strategies.

## Introduction

Sleep apnea (SA) is considered the second most frequent breathing disorder worldwide, being surpassed only by asthma, with nearly 1 billion adults between the age of 30 to 69 years suffering from mild to severe SA (1). The rapid successions of intermittent hypoxia (IH) episodes, resulting from the repetitive closing of the upper airways, contribute to the establishment and progression of several cardiovascular, respiratory, metabolic, and/or neurological comorbidities (2, 3). Recent experimental data obtained on rodent models have established that IH also alters lung functions (4–9), supporting clinical data and showing close associations between sleep apneas and lung diseases such as pulmonary arterial hypertension (PAH), asthma, chronic obstructive pulmonary disease (COPD) and fibrosis (7, 10–12).

In mice exposed to IH for 9 days, single cell RNA sequencing (scRNA-seq) of the lungs have revealed an expression profile closely resembling that observed in patients with pulmonary diseases such as PAH, COPD, and asthma, along with an important positive enrichment in pathways related to circadian clocks and energy metabolism in several cell types, and negative enrichments of pathways involved in immune responses (13). Although highly informative, this previous study was not designed to elucidate the dynamic transcriptional shifts occurring while lung cells are responding to the hypoxic challenge but rather captured a delayed snapshot of gene expression profiles several hours after the hypoxic episodes. This is a common design in sleep apnea models, as rodents are exposed to daytime IH, coinciding with their normal sleep or inactive phase, and, for convenient reasons, experiments or tissue sampling are mostly performed early morning after mice have been exposed to normoxia overnight (wake or active phase of nocturnal rodents).

In the present study, we exposed adult male mice to normoxia (Nx) or daytime IH (6:00 am to 6:00 pm) for 14 days and used samples from mice that were anesthetized while being removed from the hypoxic chamber (at 9:00 am) immediately after a hypoxic episode. We used the lungs pooled from 3 mice/groups to generate scRNA-seq data and capture cellular heterogeneity under a specific IH stimulus. Using the Azimuth Mouse Lung CellRef web interface, we identified 33 different transcription profile clusters. We performed analysis using a pseudo-bulk approach to identify the most important differentially expressed genes and enriched pathways and subsequently analyzed individual cells. Our results identify the endothelial (lymphatic and arterial) and airway smooth muscle cells as being the strongest responders to IH in terms of number of up or downregulated genes, with strong enrichments indicating myogenesis and epithelial to mesenchymal transition, as well as metabolic responses related to mitochondrial oxidative phosphorylation. A closer look at the alveolar-capillary unit cells (alveolar type 1 and 2 cells, capillary 1 and 2, alveolar fibroblast 1 and 2 and alveolar macrophages) revealed a coordinated mitochondrial remodeling across these cells in response to IH with enhanced expression of nuclear-encoded genes contributing to oxidative phosphorylation. Selective downregulation of mitochondrial encoded genes was also apparent in endothelial cells (lymphatic and arterial) and alveolar-capillary unit cells, showing mito-nuclear imbalance. We conclude that lung cells undergo specific metabolic remodeling during exposure to IH. Unsurprisingly, these responses appear to differ from those reported during the recovery phase, several hours after the last bouts of hypoxia (13). Elucidating how cells cycle between these different phases (immediate IH response vs recovery) might help understanding the various comorbidities in SA patients.

## Material and methods

### Animal and ethical approval

All experiments have been approved by the animal protection committee of Université Laval in accordance with the Canadian Council on Animal Care in Science (project# VRR-24-1515). We used a total of 6 C57BL/6NCrl male mice ordered from Charles-River Laboratories (Saint-Constant, QC, Canada) at 8-12 weeks of age. All mice were provided food and water ad-libitum and were maintained on a 12/12 h light/dark cycle. All mice were acclimatized to their new environment for 4 weeks after their arrival at our animal house facility and then randomly assigned to the IH or Nx group.

### Intermittent hypoxia

Mice were housed in standard cages (3 mice/cage) connected to an oxycycler (Biospherix, Redfield, NY, USA). Oxygen dropped from 21% to 6% in 75 seconds then returned to 21% in 45 seconds, for a total cycle length of 120 seconds and a frequency of 30 cycles/h. Mice were exposed daily for 14 days between 6:00 am and 6:00 pm. On the last day of exposure mice were euthanized at 9:00 am, immediately following a hypoxic episode, using an overdose of anesthetics (ketamine/xylazine). Control mice were kept in normoxia throughout the experiment. The lungs were rapidly dissected and digested to use in single-cell RNA sequencing (scRNA-seq).

### Sample preparation and single cell RNA sequencing

Immediately after dissection, the lungs were weighted, then minced mechanically and pooled (3 mice for each group) before digestion with cell lysis buffer supplemented with protease (Pronase, Elastase, Dispase, and Collagenase) and DNAse as described previously (14). Dissociated cells were then filtered, washed, and counted for viability. The Chromium Single Cell technology platform (10x Genomics) was used for scRNA-seq downstream processing. Briefly, for droplets generation, a total of 9100 and 7800 cells per group (Nx and IH respectively) were loaded into a Chromium Next GEM chip G (10X Genomics, cat# 1000127) and partitioned into droplets with oil and gel beads using the Chromium Controller. After emulsions were formed, barcoded reverse transcription of RNA was performed followed by libraries preparation as per manufacturer’s protocol (10x Genomics Chromium Next GEM Single Cell 3’ Library Kit v3.1, cat# 1000128). cDNA libraries were then quantified, pooled and sequenced following manufacturer’s recommendations on the Illumina NextSeq 2000 system. The total number of reads was ∼213 and 262 million with a 51 and 62% saturation at a mean depth of ∼23,000 and 33,000 reads/cell for the Nx and IH group respectively.

### Single cell data processing

We performed demultiplexing, alignment and transcript counting using Cell Ranger software (version 7.1.0) to create the gene expression matrices. The Genome Reference Consortium Mouse Built 38 (GRCm38) was used as the Mus musculus reference transcriptome for gene identification. After carrying out preliminary quality control filtration using Cell Ranger software, we obtained the mRNA sequences for 7115 Nx cells and 6179 IH cells. Bioinformatic procedures and R software (version 4.5.0) were then used to upload the matrices in Seurat (version 5.3.0) - (15) to create an object (version 5.1.0) and filter matrices for further quality controls as well as to cluster, visualize, and annotate cells. Another round of quality control using nCount (number of UMI – unique molecular identifiers – per cell), nFeature_RNA (number of genes expressed per cell) and mt_percent (fraction of UMIs corresponding to mitochondrial genes) was applied to filter low quality cells. Thresholds to identify outliers were calculated using the isOutlier function from the scater package (version 1.36.0) as nCount fewer than 22970 and 17131 UMI, nFeature fewer than 7038 and 6181 expressed genes, and percent.mt under 6.31 and 4.93 percent (for IH and Nx-cells respectively, a table of the applied filters can be seen in supplemental table 1). Data were normalized using SCTransform (version 0.4.2), then we identified and filtered doublets (when two or more cells are captured into a single oil droplet) using the R library DoubletFinder (version 2.0.6) and assuming a 5% doublet rate. Cells were clustered based on their transcription profile using the Azimuth Mouse Lung CellRef (16). We generated violin plots of quality control metrics and dimensionality reduction visualizations using UMAP of predicted cell lineages at levels 1 and 3 using the ggplot2 package (version 3.5.2 – Figure 2 A-C). The filtered and normalized scRNAseq data were utilized to identify differentially expressed genes across predicted cell types. Differential expression analysis was performed using the FindAllMarkers function, with cells grouped according to their predicted identities. For each cluster, the top 20 most specific marker genes were selected based on their expression profiles. These markers were subsequently visualized using a heatmap generated with the DoHeatmap function (Figure 2 D).

### Cell type proportions

Since tissues from three animals per group were pooled (n = 1 per condition), standard statistical tests were not applicable. Accordingly, we performed an exploratory analysis using a bootstrap-based approach to compare cell type proportions between IH and Nx. We resampled cells with replacement to estimate the distribution of each cell type’s proportion in IH and Nx and calculated 95% confidence intervals for their differences. Although this method assumes cell-level independence (a simplification, as cells come from the same animals), it leverages the large number of cells to estimate variability in the absence of biological replicates. To assess whether observed differences could arise by chance, we also performed a permutation test: condition labels (IH vs Nx) were randomly shuffled 1000 times, and the absolute difference in cell counts recalculated each time. Empirical p-values were computed as the proportion of permutations with a difference equal to or greater than the observed one (with a continuity correction).

### Gene expression and GSEA analyses

Downward analyses and graphical data presentations were done using the BigOmics platform (https://bigomics.ch – Omic Playground v3.5.24). Because the platform cannot accommodate more than 2000 cells per datasets, cells were aggregated using the Supercell.R package (17). The SuperCell algorithm parameters were set at an upper limit of 2000, which generated 1793 metacells (935 IH and 858 Nx) from our initial total of 13291 cells (6179 IH and 7115 Nx). We compared the cell type distribution and proportions between the original single cell and the metacells (supplemental Figure 1), showing a high Pearson’s correlation coefficient (R=0.9, p<0.0001 – supplemental Figure 1B). Thus, we are confident that the metacells retained the relative abundance of cell types during aggregation.

We used the platform to evaluate differentially expressed genes (DEG), compute the number of up- and down-regulated genes, draw the volcano plots, genes and gene sets UMAP, and perform Gene Set Enrichment Analysis (GSEA). Up - and down-regulated genes in response to IH were discriminated by using the linear model for microarray data (limma) trend test rather than simple t-test or Welch’s t-test to gain more power related to the empirical Bayes moderation and increased degree of freedom of this test. For all volcano plots, DEG were filtered for a false discovery rate (FDR) < 0.05 and a log2 fold change (log2FC) > 0.5 or < -0.5. Gene Set Enrichment Analysis (GSEA) was performed using the Hallmark collection of gene sets from MSigDB, which aggregate and refine overlapping MSigDB gene sets into 50 high-level non-redundant biological themes (18). Both fgsea and Fisher’s exact test were applied (with an FDR threshold <0.1) and sorted by normalized enrichment scores (NES) to highlight the most significant pathways.

### Data availability

All data used in the present analysis are publicly available through the NCBI Gene Expression Omnibus with the GEO accession number GSE301350.

## Results

### scRNA sequencing and cell identification following exposure to intermittent hypoxia

A visual representation of the methods can be seen in Figure 1. Violin plots of global quality of the cells before downstream analysis show that the distribution of detected genes (nFeature_RNA) and transcript counts (nCount_RNA) are similar across conditions (Figure 2A), indicating comparable sequencing depth and complexity. The proportion of mitochondrial genes (percent.mt) remains largely under 20%, suggesting minimal stress or apoptotic contamination. Using the LungMap single-cell reference Azimuth Mouse Lung CellRef web interface, predicted lineage level 1 annotation revealed 4 general clusters: epithelial (≈7%), endothelial (≈16%), mesenchymal (≈13%), and immune (≈63% - see Figure 2B for the UMAP projection). Automated cell annotation level 3 identified a total of 33 different clusters that are presented as a UMAP projection (Figure 2C) and heatmap showing gene expression level of the markers used to identify each cell type (Figure 2D). This is in line with the 40 different cell types reported by CellRefs in the lungs of mice under highly optimized sequencing depth (16). Level 3 identified 33 cell types in IH-cells versus 29 in Nx-cells (Table 1). B cells were expressed in the largest proportion (25% and 23% of total cell count in Nx and IH animals respectively – Table 1). The total count of immune cells represented 66% and 55% in Nx- and IH-cells respectively (supplemental Table 2), while the alveolar capillary unit cell population (capillaries 1 and 2, alveolar fibroblasts 1 and 2, alveolar type 1 and 2 cells, and alveolar macrophages), represented respectively 28 and 43% of the total cell count in Nx- and IH-cells (supplemental Table 2). The bootstrap estimated differences of cell type proportion are reported in supplemental Figure 2, suggesting that IH increases the number of AM, AT1, AF2 and Cap1 cells, and decreases the number of Neutrophil, iMON and LEC (see abbreviations for all cell types in Table 1) which are the cell types showing the most robust changes. However, the reported differences where no longer significant after the permutation testing, and the results should therefore be interpreted with caution.

**Figure 1:**
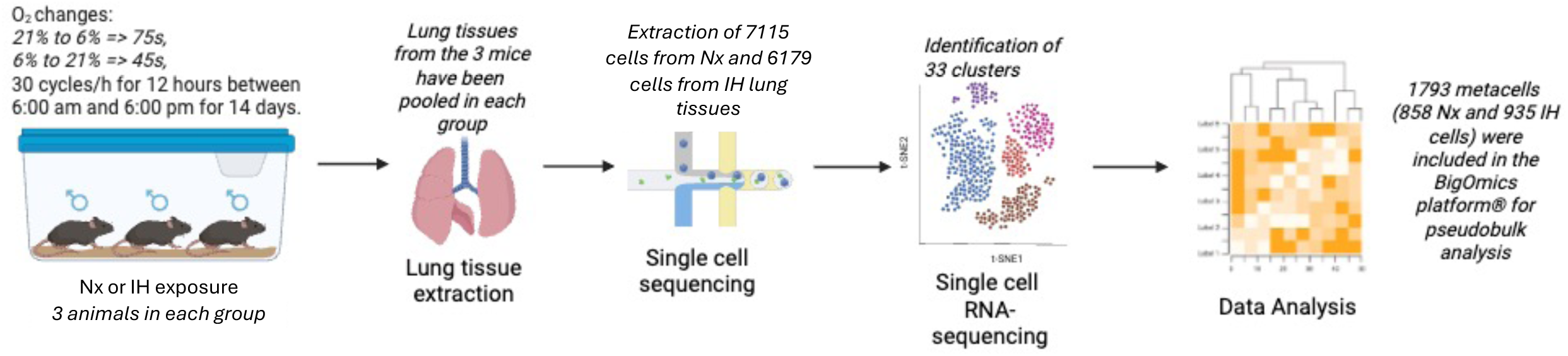
Experimental design. Adult male mice were exposed to room air (Nx – 21% O_2_) or intermittent hypoxia (IH – 21 to 6% O_2_) for two weeks, a total of 6 lungs were harvested (3 per group) and digested for scRNA sequencing and downstream analysis. Generated using BioRender.

**Figure 2:**
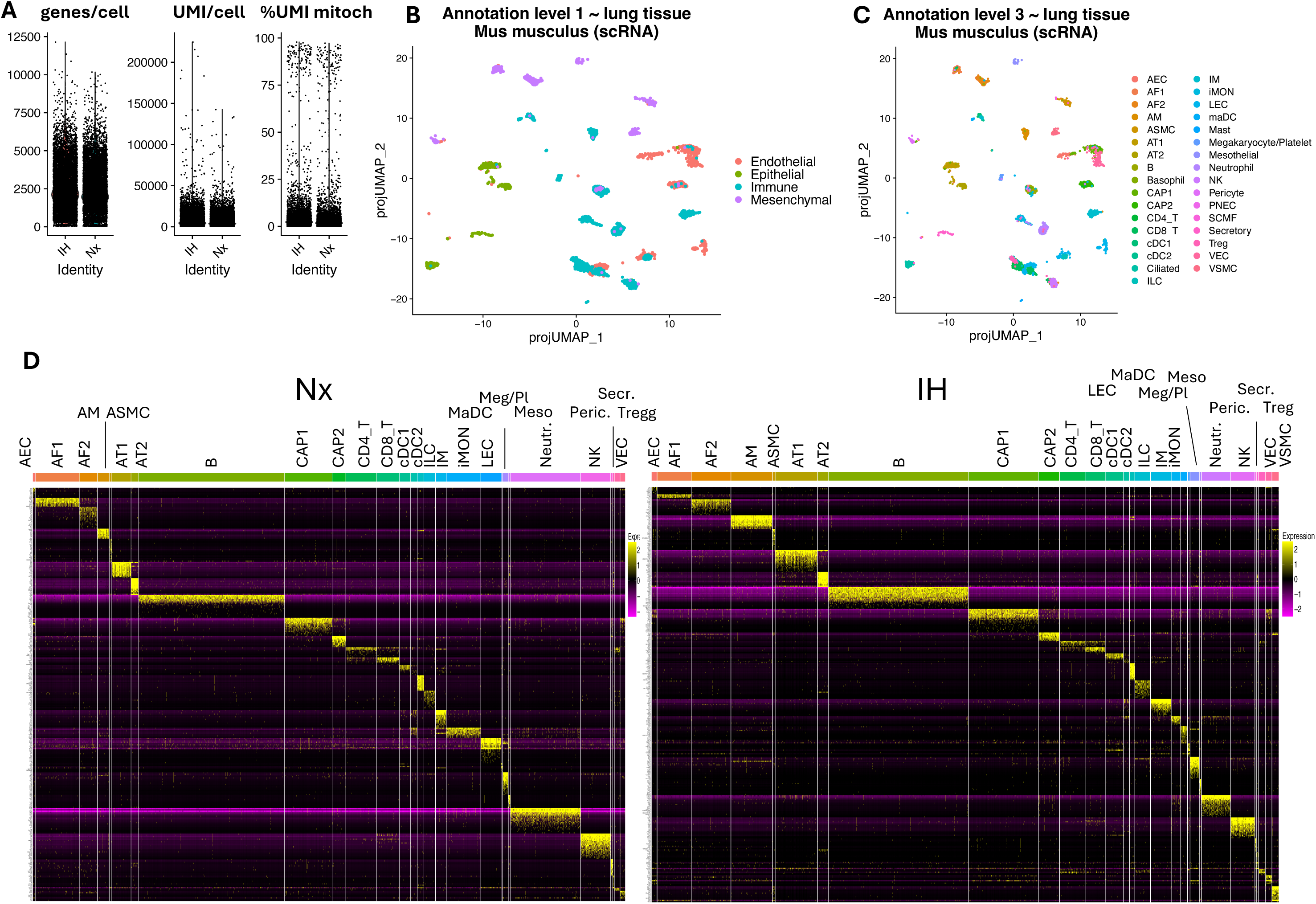
Cell quality, Azimuth-based cell annotation, and marker gene expression in single-cell RNA-sequencing (scRNA-seq). (A) Violin plot representation of global quality of the cells before downstream analysis, representing nFeature (#genes/cell), nCount (#UMI/cell) and percent.mt (%UMI of mitochondrial genes). (B and C) UMAP plots of all scRNA-seq data showing the general (level 1 annotation – B) and specific (level 3 annotation – C) cell types that were identified. (D) Heatmap of the gene expression level of marker genes in each identified cluster/cell type in mice exposed to Nx (left) or IH (right).

**Table 1:** Identified cell population and proportions at levels 1 and 3 in mice exposed to normoxia (Nx) or intermittent hypoxia (IH). Endo. : Endothelial cells. Epi. : Epithelial cells. Immun. : Immune cells. Mes. : Mesenchymal cells.

| Identified CellCards population |  |  | Number of cells |  | Proportion % |  |
| --- | --- | --- | --- | --- | --- | --- |
| abbreviation | Level 1 cell group | Level 3 cell type | Nx | IH | Nx | IH |
| AEC | Endo. | arterial endothelial cells | 33 | 58 | 0,5% | 0,9% |
| CAP1 | Endo. | capillary 1 cells | 575 | 696 | 8,1% | 11,3% |
| CAP2 | Endo. | capillary 2 cells | 164 | 207 | 2,3% | 3,4% |
| LEC | Endo. | lymphatic endothelial cell | 241 | 64 | 3,4% | 1,0% |
| VEC | Endo. | venous endothelial cells | 61 | 61 | 0,9% | 1,0% |
| AT1 | Epi. | alveolar type 1 cells | 219 | 415 | 3,1% | 6,7% |
| AT2 | Epi. | alveolar type 2 cells | 86 | 103 | 1,2% | 1,7% |
| Ciliated | Epi. | ciliated cells | 75 | 48 | 1,1% | 0,8% |
| PNEC | Epi. | pulmonary neuroendocrine cells | 0 | 2 | 0,0% | 0,0% |
| Secretory | Epi. | secretory cells | 13 | 18 | 0,2% | 0,3% |
| AM | Immun. | alveolar macrophages | 138 | 413 | 1,9% | 6,7% |
| B | Immun. | B cells | 1782 | 1405 | 25,0% | 22,7% |
| Basophil | Immun. | basophils | 1 | 1 | 0,0% | 0,0% |
| CD4_T | Immun. | CD4 T cells | 377 | 252 | 5,3% | 4,1% |
| CD8_T | Immun. | CD8 T cells | 266 | 196 | 3,7% | 3,2% |
| cDC1 | Immun. | dendritic cells 1 | 134 | 180 | 1,9% | 2,9% |
| cDC2 | Immun. | dendritic cells 2 | 78 | 59 | 1,1% | 1,0% |
| ILC | Immun. | innate lymphoid cells | 137 | 151 | 1,9% | 2,4% |
| IM | Immun. | interstitial macrophages | 127 | 165 | 1,8% | 2,7% |
| iMON | Immun. | inflammatory monocytes | 421 | 86 | 5,9% | 1,4% |
| maDC | Immun. | dendritic cells | 12 | 20 | 0,2% | 0,3% |
| Mast | Immun. | mast cells | 0 | 2 | 0,0% | 0,0% |
| Megakaryocyte/<br>Platelet | Immun. | megakaryocyte/platelet | 46 | 89 | 0,6% | 1,4% |
| Neutrophil | Immun. | neutrophils | 848 | 271 | 11,9% | 4,4% |
| NK | Immun. | natural killer cells | 365 | 240 | 5,1% | 3,9% |
| Treg | Immun. | regulatory T cells | 66 | 62 | 0,9% | 1,0% |
| AF1 | Mes. | alveolar fibroblast 1 cells | 540 | 366 | 7,6% | 5,9% |
| AF2 | Mes. | alveolar fibroblast 2 cells | 264 | 443 | 3,7% | 7,2% |
| ASMC | Mes. | airway smooth muscle cells | 25 | 25 | 0,4% | 0,4% |
| Mesothelial | Mes. | mesothelial cells | 8 | 13 | 0,1% | 0,2% |
| Pericyte | Mes. | pericytes | 13 | 20 | 0,2% | 0,3% |
| SCMF | Mes. | secondary crest myofibroblasts | 0 | 2 | 0,0% | 0,0% |
| VSMC | Mes. | vascular smooth muscle cells | 0 | 43 | 0,0% | 0,7% |
| Total number of cells present in the lungs |  |  | 7115 | 6176 | 100,0% | 100,0% |

### Gene expression and pathway enrichments in response to IH in the pseudo-bulk analysis

The UMAP projection of DEG from the pseudo bulk analysis shows clear clustering of up (red dots) and down (blue dots) regulated genes (Figure 3A). When filtered for significance (FDR < 0.05) and effect size (log2FC > +/- 0.5), there were 837 DEG in IH vs Nx, out of which 574 were upregulated and 263 were downregulated (Figure 3B and supplemental data 3B). Among the 10 most up regulated genes in response to IH 6 were nuclear-encoded mitochondrial genes, including subunits of electron transport chain complex I (Ndufa13, Ndufb11, Ndufb7), ATP synthase (Atp5g1, Atp5d), and a component of the mitochondrial translocase allowing protein transport from the cytosol (Tomm6), suggesting a mitochondrial response to IH (Figure 3B). A visual examination of the UMAP projection showed that these genes are clustered in the small box highlighted in panel 3A. Conversely, mitochondrial genome-encoded genes, mt-Nd6 and mt-Atp8, were among the top 10 downregulated genes, indicating a potential mito-nuclear gene expression imbalance. The gene set UMAP (Figure 3C) also showed strong clustering of pathways (all collections of gene sets present in the Bigomics platform confounded). Restricting the gene sets to Hallmark collections (Figure 3D and supplemental data 3D), GSEA confirmed upregulated responses pertaining to core components of cellular respiration and mitochondrial energy metabolism, including oxidative phosphorylation, and fatty acid metabolism, but also reactive oxygen species (ROS), the MYC targets V1 (a protooncogene pathway), DNA repair, myogenesis, peroxisome and the Epithelial Mesenchymal transition. The negatively enriched pathways included processes related to inflammatory responses, cell cycle (Mitotic spindle) and cell cycle check points, but also androgen response, which appears as a relevant information since we previously reported interactions between testosterone levels and pulmonary responses to IH (4).

**Figure 3:**
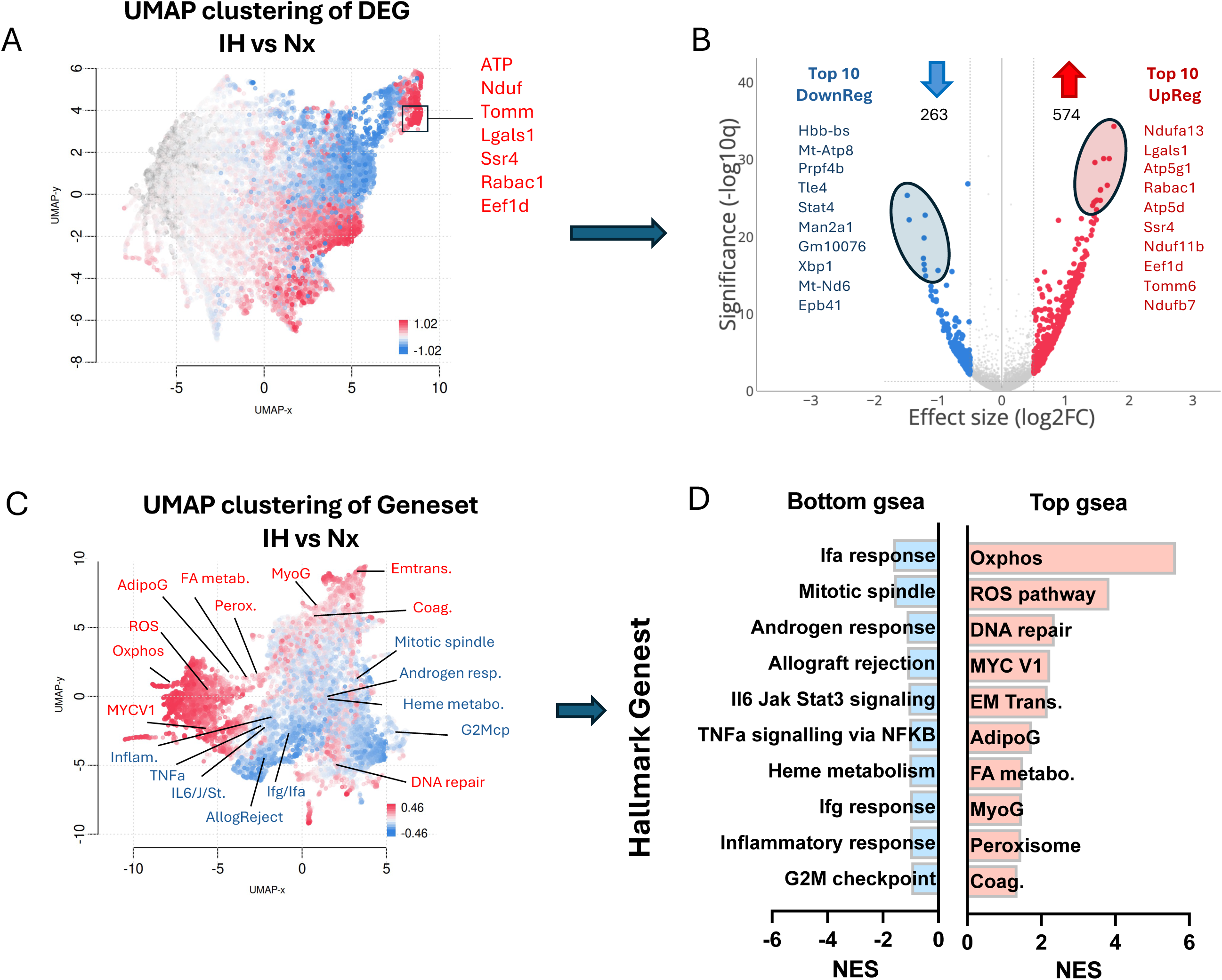
UMAP clustering in a pseudobulk analysis of gene expression and pathway enrichment in mice exposed to Nx or IH. (A) UMAP clustering of differentially expressed genes (DEG – red: upregulated, blue: downregulated). (B) Volcano plot of DEG and identity of the 10 most upregulated and top 10 most downregulated genes. Significant thresholds for volcano plots: q value < 0.05 and log2FC > +/- 0.5. (C) UMAP clustering of pathways (gene sets) and localization of the top 10 positively (red) and negatively (blue) enriched pathways. (D) Top 10 positively and negatively enriched pathways in the Hallmark collection of gene sets. NES: normalized enrichment score.

### Cell-specific pathway enrichments in response to IH

We extended the GSEA analysis (restricted to the Hallmark collection) to the 28 cell types identified both in IH and Nx groups. By focussing on the top 20 enriched gene sets, the analysis was restricted to 20 cell types (Figure 4). Oxidative phosphorylation was significantly enriched with high NES in most cell types present in the lungs, the only exception being the AEC, AF1, ASMC, and mesothelial cells (Msth). Other pathways showing enrichments in several cell types include MYC Targets V1 (Cap1, CD8, cDC2, IM, LEC, Msth, and regulatory T cells), DNA repair (AF2, B, cDC2, IM, LEC, Scrt, Treg, and VEC), ROS pathway (AEC, AT1, B, NK), and epithelial-mesenchymal transition (ASMC, AT1, LEC, and pericytes). Before going into further details regarding these pathways, we assessed the intensity of the transcriptomic responses at the single cell resolution by computing the number of differentially expressed genes within each cell types.

**Figure 4:**
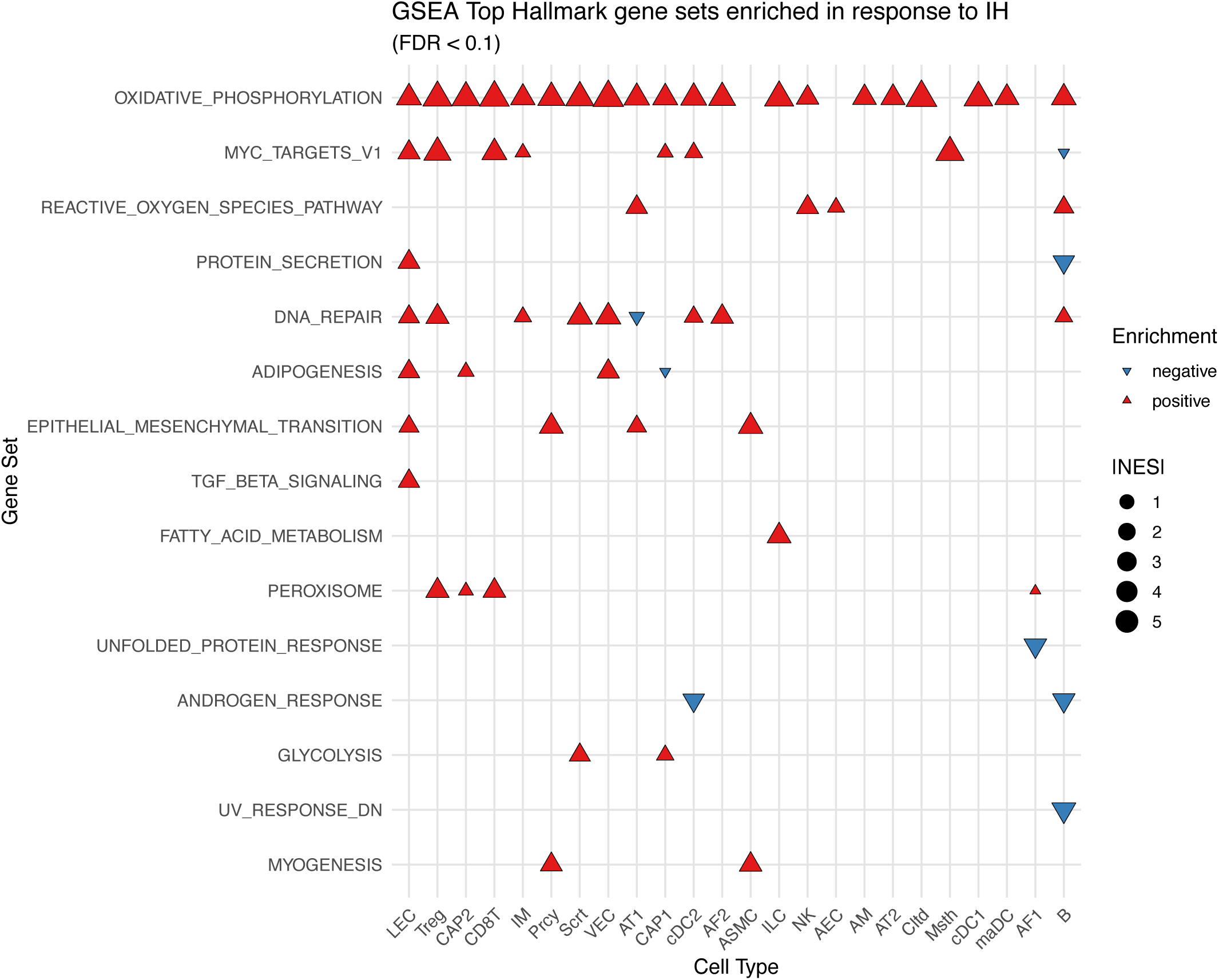
Gene set enrichment analysis across cell types in mice exposed to Nx or IH. Table of the 20 most enriched pathways ranked using highest values of rms.NES (top to bottom), then, cell types were organized according to count of positive and negative enrichments (highest score on the left). |NES| : absolute value of the normalized enrichment score. Significant threshold: FDR<0.1

### IH induced profound transcriptional reprogramming in ASMC, LEC and AEC

Figure 5A reveals the cell type specificities of the response to IH in the lungs. Airway smooth muscle cells (ASMC) and lymphatic endothelial cells (LEC) predominantly exhibit upregulation of gene expression in IH (1385 and 1324 upregulated genes respectively). Intriguingly, LEC show a pronounced activation of gene transcription despite having a much lower number of cells under IH compared with Nx (64 vs 241 respectively). Conversely, arterial endothelial cells (AEC) show a predominant downregulation of gene expression in IH (879 downregulated genes). Other cell populations, show varying degree in the number of genes up or down regulated, and a few, such as cCD1, CAP2, CD8-T, NK, IM and B show less than 100 genes affected by IH.

**Figure 5:**
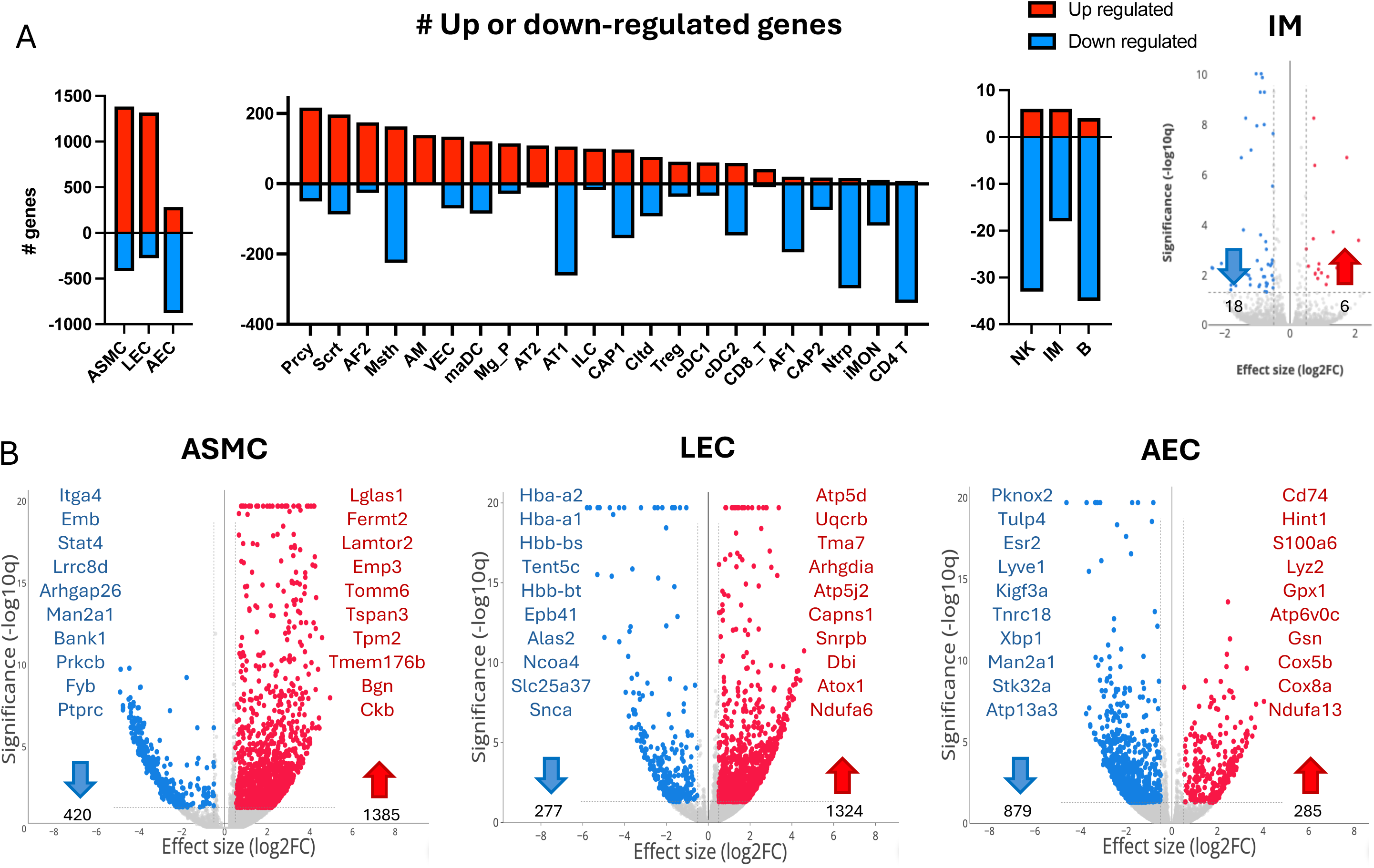
IH induced profound transcriptional reprogramming in ASMC, LEC and AEC. (A) Number of significantly altered genes in the lung cells when comparing IH to Nx. (B) Volcano plots showing the up- (red dots) and down-regulated (blue dots) genes in response to IH exposure in ASMC, LEC, and AEC. Top 10 up- and down-regulated genes are listed beside the volcano. The volcano plot for IM (upper right corner), illustrates a cell type with minimal response to IH (see gene list in supplemental data 5B). (In all panel, statistical methods are trend.limma and significant thresholds: q value < 0.05, logFC > ±0.5).

Focusing on ASMC, LEC and AEC we analyzed individual differential gene expression and performed GSEA using the Hallmark collection of gene sets. In ASMC the 10 most upregulated genes (Figure 5B and supplemental data 5B) included galectin-1 (Lgals1) which prevents pathological vascular remodeling and regulates mitochondrial function (19), and biglycan (Bgn) and tropomyosin beta chain (Tpm2) which are involved in extracellular matrix and contractile functions. Mitochondrial response in ASMC is limited to the increased expression of Tomm6 (component of mitochondrial protein import), while the increased expression of creatine kinase (Ckb), which can generate ATP rapidly in response to an abrupt increase in metabolic demand without engaging glycolysis or oxidative phosphorylation, supports the idea that energy buffering mechanisms are upregulated in IH. The 10 most downregulated genes pertain to immune and cytoskeletal signaling pathways, also consistent with airway remodeling.

In LECs, results showed the upregulation of multiple nuclear-encoded mitochondrial genes: Atp5d, Uqcrb, Atp5j2, Ndufa6, and the chaperone Atox1 that supports antioxidant defenses by binding free metals in endothelium (20), suggesting that LECs exposed to IH are actively remodeling their mitochondrial machinery. LEC’s 10 most downregulated genes included genes involved in iron homeostasis and heme synthesis (several related to hemoglobin), including Slc25a37 (a mitochondrial iron transporter), Ncoa4, and Alas2 (Figure 5B, supplemental data 5B).

In AECs, X-box binding protein 1 (Xbp1), one of the 10-most downregulated genes in response to IH, is a key transcription factor involved in the unfolded protein response that plays a crucial role in regulating mitochondrial function and morphology, particularly in response to endoplasmic reticulum stress, and can also deeply modify mitochondrial ROS production and mitophagy (21). Downregulation of Xbp1 may reflect impaired adaptive responses to proteotoxic stress, which can secondarily affect mitochondrial health. The remaining downregulated genes (Atp13a3, Stk32a, Man2a1, Tnrc18, Kif13a, Lyve1, Esr2, Tulp4, and Pknox2) are mostly involved in diverse cellular processes such as cell signaling (including estradiol signaling through the Estradiol receptor β - Esr2) transport, or differentiation. Regarding upregulated genes in AEC, results showed the upregulation of key mitochondrial and antioxidant genes. Indeed, the enhanced expression of Gpx1, Cox5b, Cox8a, and Ndufa13 suggests that AECs are potentially increasing their electron transport chain capacity and antioxidant defenses (Figure 5B and supplemental data 5B). The volcano plot for IM (Figure 5), illustrates a cell type with minimal response to IH (see gene list in supplemental data 5B).

We next visualized ASMC, LEC and AEC enrichment scores on UMAPs by summarizing all available collections of gene sets (Figure 6 A-C). This revealed broadly positive enrichment in ASMC and LEC and negative enrichment in AEC. We then performed targeted GSEA using the Hallmark collection of gene sets (Figure 6D and supplemental data 6D). In ASMC, epithelial-mesenchymal transition (EMT – known to be activated under hypoxic conditions (22)) and myogenesis were enriched, suggesting that phenotypes associated to the development of PAH, lung fibrosis and asthma (23, 24) are induced during IH. While oxidative phosphorylation had a high NES in ASMC, the q value (0.4) was above the significance threshold of 0.1. In LEC, several pathways were significantly enriched, also including EMT and TGF beta signaling (one of the positive regulators of the EMT, that can be activated by ROS and hypoxia (22)), as well as oxidative phosphorylation, MYC target V1 (an oncogene pathway), and DNA repair (suggesting damage to DNA are induced by IH). In AEC, only the ROS pathway was significantly enriched, the oxidative phosphorylation pathway showed a strong normalized enrichment score (NES = 2.79) but did not strictly meet the FDR<0.1 threshold (q=0.102), this might reflect limited statistical power when pooling 3 animals per conditions. Genes composing the Hallmark ROS gene set included Atox 1 (that supports antioxidant defenses by binding free metals), glutathione peroxidase 4 (Gpx4 – an essential antioxidant enzyme), and other antioxidant such as thioredoxin 1 and peroxiredoxin 6, that can have antioxidant peroxidase activity in the lungs but also prooxidant effects through NOX2 (25).

**Figure 6:**
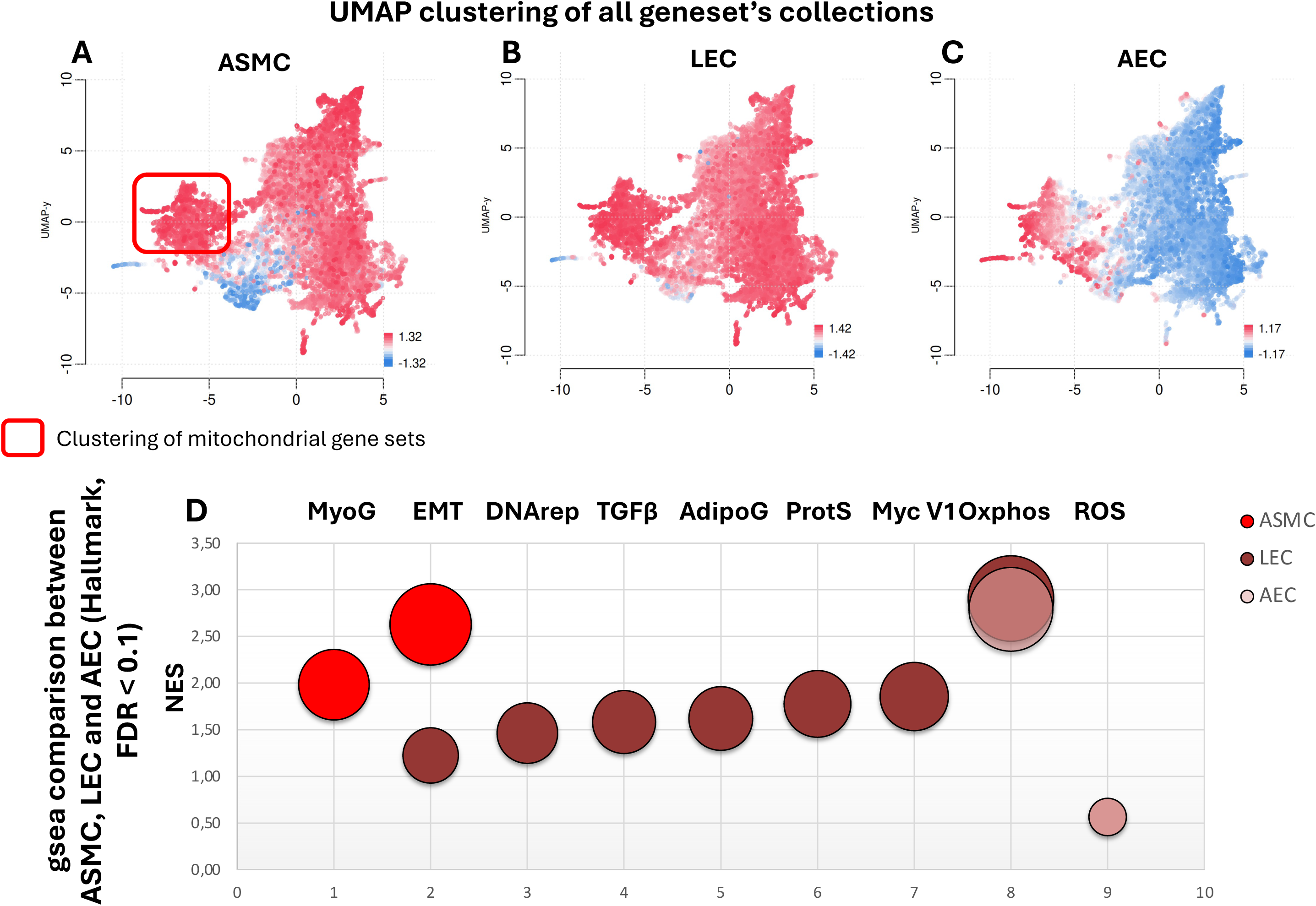
Gene set enrichment analysis in ASMC, LEC and AEC in mice exposed to Nx or IH. (A-C) UMAP clustering of pathways (gene sets) in ASMC, LEC and AEC. (D) Bubble plot of significantly enriched pathways (Hallmark collection of gene sets - FDR<0.1) in ASMC, LEC and AEC.

It is noteworthy that pathways related to mitochondrial respiration are clustered in the red highlighted box in Figure 6 A (Box selection returned 1557 gene sets including 123 gene sets from the gene ontology collection – see supplemental data 6A). Targeted GSEA using the gene ontology (GO) collection in ASMC, LEC and AEC cells, showed a large variety of pathways related to mitochondrial electron transport chain and antioxidant peroxidase activity in AEC and LEC but not ASMC (see supplemental Figure 3 and supplemental data 6A-D). Furthermore, the negative enrichment of 3 pathways related to the transforming growth factor β in the GO_BP collection (not apparent in the Hallmark collection) in AEC suggest responses involving a key pathway determinant of pulmonary health (26) (see supplemental Figure 3 and supplemental data 6A-D).

### The specificity of the alveolar-capillary unit cells

We further looked into the specificity of the alveolar epithelium-capillary interface, responsible for respiratory gas exchange, by including the alveolar type 1 and 2 cells (AT1 and AT2), alveolar macrophages (AM), the capillary endothelial cell types 1 and 2 (CAP1 and CAP2) and the alveolar fibroblasts type 1 and 2 (AF1 and AF2, Figure 7A) (27). As noted above, cell counts of the alveolar-capillary unit represented 28% and 43% of total Nx- and IH-cells respectively (see supplemental Table 2). Under IH, AT1 cells roughly doubled in number (219 Nx-cells vs 415 IH-cells – Table 1), while AT2 cells, which serve as facultative progenitors for AT1, increased more modestly (86 Nx-cells vs 103 IH-cells). The number of AM raised sharply (138 Nx-cells vs 413 IH-cells), pointing to enhanced immune surveillance, debris clearance or remodeling signals that often accompany epithelial repair (28, 29). In parallel, we saw a relative decrease in quiescent fibroblasts (AF1) and an increase in activated-like fibroblasts (AF2), hinting at extracellular-matrix deposition (30).

**Figure 7:**
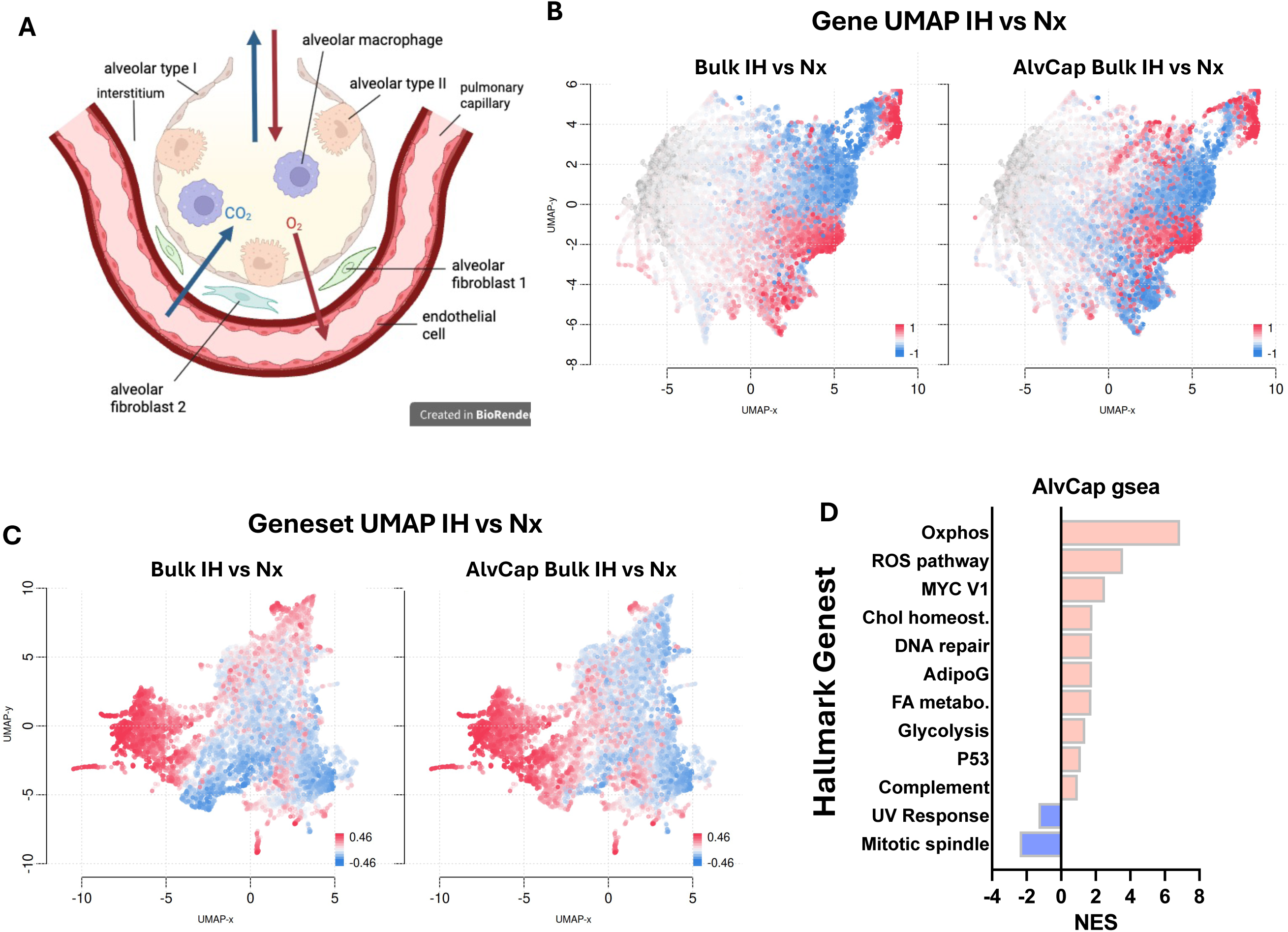
Response of the alveolar capillary unit in mice exposed to Nx or IH. (A) Depiction of cells forming the alveolar capillary (AlvCap) unit. (B) UMAP clustering of differentially expressed genes (DEG – red: upregulated, blue: downregulated) in the pseudobulk whole lung data set (left) and the AlvCap data set (right). (C) UMAP clustering of positively (red) and negatively (blue) enriched pathways in the pseudobulk whole lung data set (left) and the AlvCap data set (right). (D) Top 10 positively and 2 negatively enriched pathways in the Hallmark collection of gene sets. NES: normalized enrichment score. Significant threshold : FDR<0.1

We used a bulk approach to compare the pseudo-‘AlvCap Bulk’ (where we pooled only cells within the alveolar-capillary unit) with the whole lung pseudo bulk. The visual comparison of gene and gene set UMAPs, shows close similarities between the two pseudo bulk sets, particularly in the left regions of the gene set UMAPs (Figure 7B and 7C), containing the mitochondrial related pathways. Targeted GSEA using the Hallmark collection to compare the significantly enriched pathways (FDR < 0.1) between the pseudo-‘AlvCap Bulk’ and whole lung pseudo bulk datasets, showed 11 shared pathways in both datasets (out of 14 significant ones in the pseudo-‘AlvCap Bulk’ – supplemental Figure 3). Oxidative phosphorylation and ROS pathways rank at the very top in both datasets, with low q-values (≤ 10^-3^) and high NES (≈ 6.89–3.58 vs. ≈5.64–3.84). Further individual analysis of cell types confirmed enrichments in the mitochondrial related pathways region (Figure 8A – all collections of gene sets present in the Bigomics platform confounded, and supplemental data 7&8) and cell-specific responses with AF2, AT2 and AM having exclusively positive gene set enrichments, while AF1, AT1, Cap1 and Cap2 displaying a mixed pattern of both up and downregulated gene sets. In all cells of the alveolar capillary unit (except AF1), the highest enriched pathway is oxidative phosphorylation. TNFɑ signaling via NFκB was significantly enriched in AT2, AT1, and Cap2 cells, apoptosis in Cap2 and AT2 cells, and ROS pathway in AT1 cells (Figure 8B).

**Figure 8:**
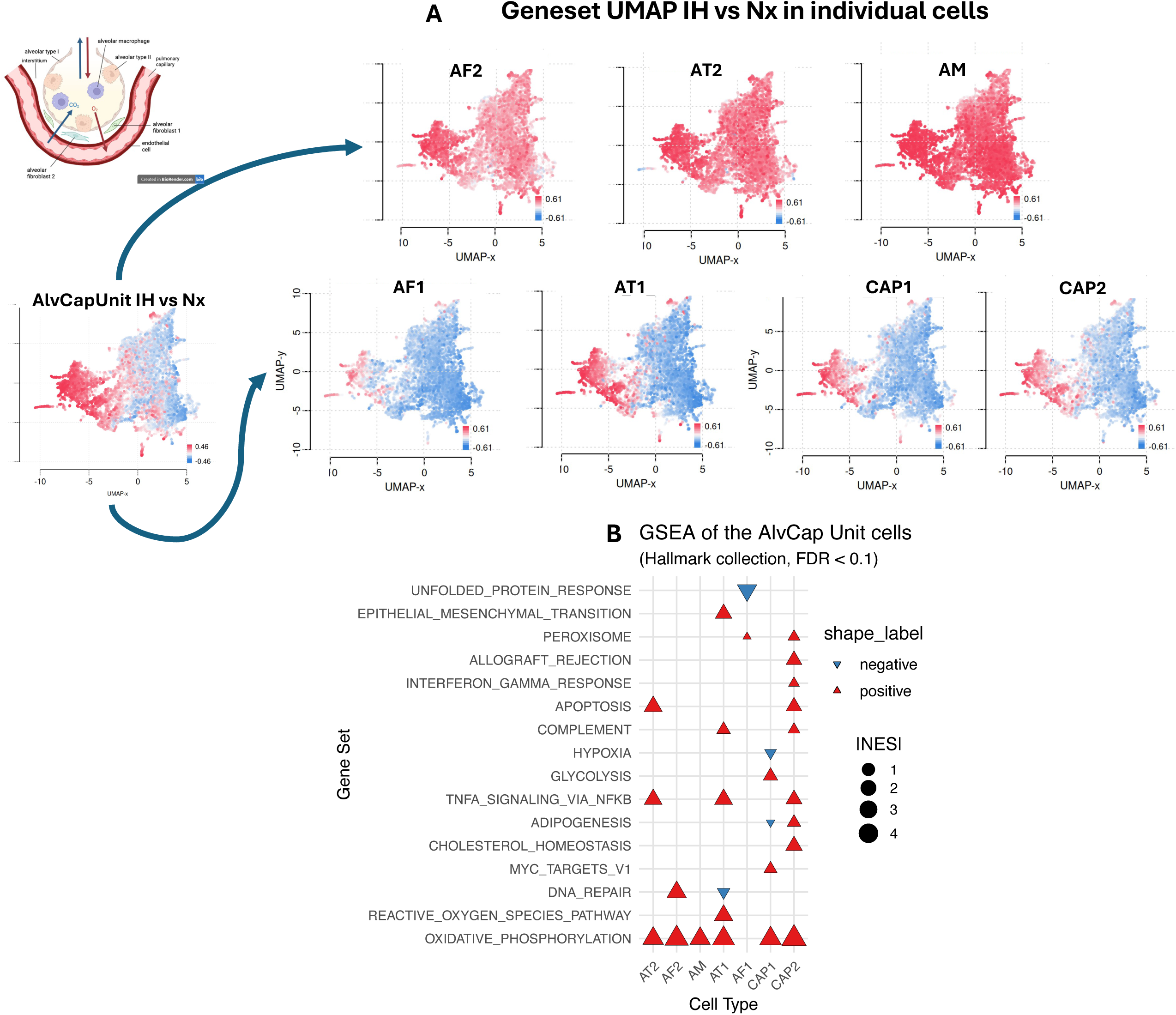
Gene set enrichment analysis of individual cells in the alveolar capillary unit in mice exposed to Nx or IH. (A) UMAP clustering of positively (red) and negatively (blue) enriched pathways at the cell level in the alveolar capillary unit. (B) Table of positively or negatively enriched pathways (Hallmark collection of gene sets). |NES| : absolute value of the normalized enrichment score. Significant threshold : FDR<0.1

The AlvCap Unit bulk data set was used to highlight the extent of upregulation of genes involved in mitochondrial oxphos (Figure 9). Most of the genes composing the nuclear-encoded subunits of Complex I (Ndufa, Ndufb, Ndufs, etc.), Complex II (SdhA-D), Complex III (Uqcrq1-2, Uqcrb, Uqcrfs1) and Complex IV (Cox5a-b, Cox6a2, Ndufa4, etc.) of the electron transport chain (ETC) are up-regulated under IH, together with the adenine nucleotide translocator slc25a4 and slc25a5 which are required for the export of ATP to the cytosol (and import of ADP). Since some mRNA from the mitochondrial DNA appeared among the top 10 downregulated genes on the lung bulk analysis (see Figure 3B) we also sought to report the expression of these genes in the diverse cell types. The data reported in supplemental Table 3 show down-regulation of 11 (out of 13) mt-genes in LEC and 6 in ASMC. In the alveolar-capillary unit AF1, AF2, AT1, Cap1 and Cap2 cells showed downregulated expression of mt-Atp8 and mt-Nd6 (AF1, AT1) or mt-Nd4l (AF2). mt-Atp8 also appears to be downregulated in a variety of immune cells. Since large differences between the expression levels of nuclear and mitochondrial genes coding for proteins of the ETC (mito-nuclear imbalance) represent a specific signature of mitochondrial stress (31–33), we sought to quantify this effect by focusing on the 67 nuclear genes of the GO_BP aerobic electron transport chain (GO:0019646) and on the 13 genes of the ETC encoded by mitochondrial DNA. For the cells showing significant enrichment of both pathways, we calculated the mean log2FC (all genes of the pathway) and reported an index of nuclear to mitochondrial imbalance as the difference mean nuc – mean mt, where a high score corelates with more pronounced mito-nuclear imbalance. The results reported in Figure 10 (and supplemental data 10) show that LEC have the strongest imbalance, followed by AEC, AF2, AT1, Cap2, Cap1 and AF1. Finally, under IH, the tricarboxylic acid (TCA) cycle itself shows a complementary pattern of transcriptional remodeling that dovetails with the changes in the oxidative-phosphorylation pathways (Figure 11). The up-regulation of TCA enzymes (IDH, Sdh, Fh, Mdh2) directly supports our GSEA and DEG findings showing enrichment of mitochondrial oxidative-phosphorylation and electron-transport pathways.

**Figure 9:**
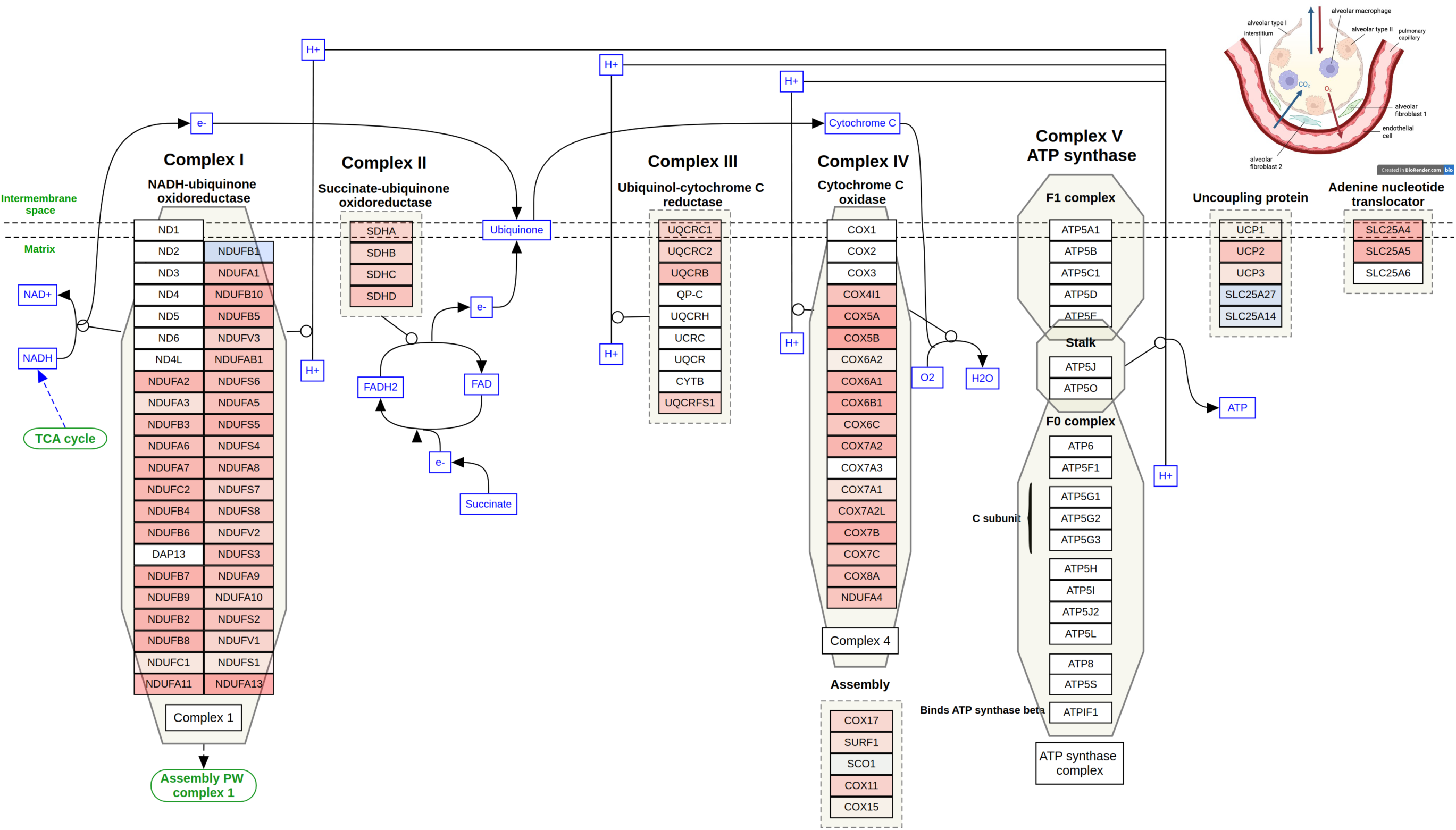
Gene expression in the electron transport chain OXPHOS system in mice exposed to Nx or IH. Genes are colored according to their upregulation (red) or downregulation (blue) in the contrast profile (pseudo-“AlvCap Bulk” IH vs Nx) on the WikiPathways_20230410_WP111_Homo sapiens.

**Figure 10:**
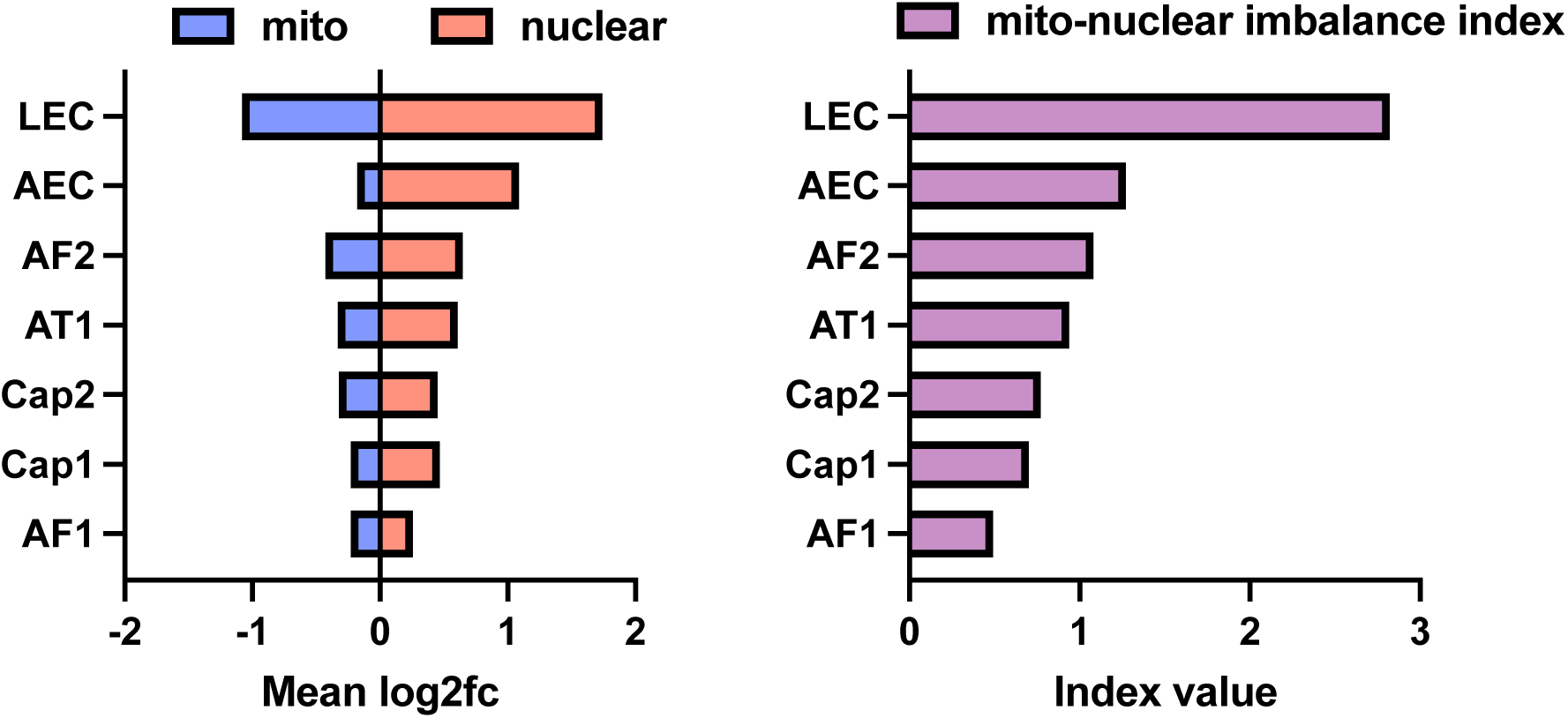
Mito-nuclear imbalance in mice exposed to Nx or IH. Mean log2FC of the nuclear DNA (GO_BP aerobic electron transport chain - GO :0019646) and electron transport chain genes from the mitochondrial DNA (left), and mito-nuclear imbalance index (nuclear mean log2FC – mitochondrial log2FC - right).

**Figure 11:**
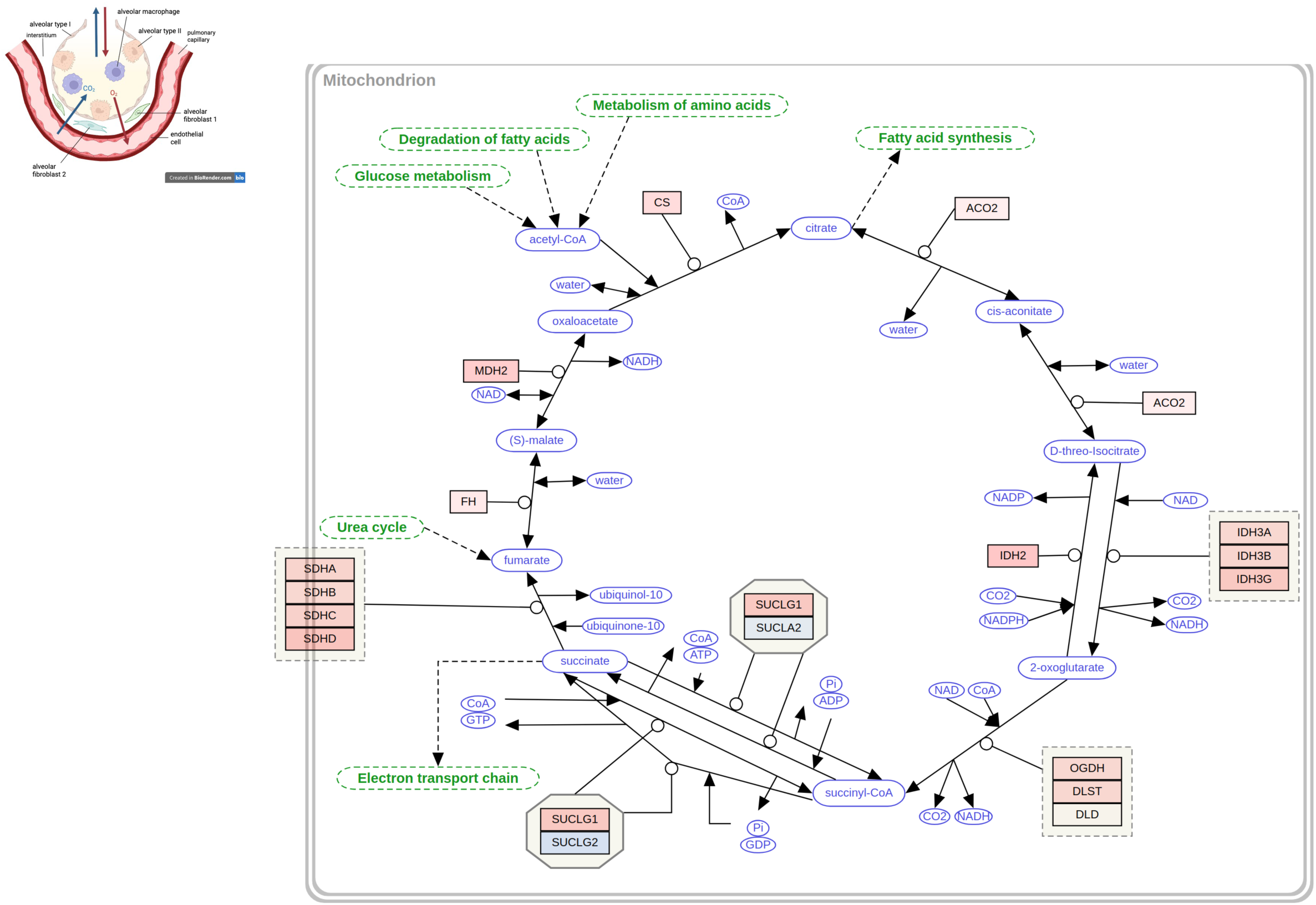
Pathway map of the TCA cycle. (WikiPathways_20230410_WP78_Homo sapiens). Genes are colored according to their upregulation (red) or downregulation (blue) in the contrast profile (pseudo-“AlvCap Bulk” IH vs Nx).

## Discussion

Our data reveal some key features of the gene expression pattern at the single cell resolution in the lungs during exposures to intermittent hypoxia. Several important observations emerge from this data set: **1.** The cells that show the most prominent responses, in terms of number of differentially expressed genes, are the airway smooth muscle cells, the lymphatic endothelial cells, and the arterial endothelial cells; **2.** Strong enrichments of myogenesis and epithelial to mesenchymal transition (EMT) in ASMC, and of oxidative phosphorylation (oxphos) and reactive oxygen species (ROS) pathway in AEC appear particularly relevant for pulmonary disease progression. **3.** There is an apparent selective reorganization of the alveolar-capillary unit with drastically increased number of individual AM, AF2 and AT1 cells; and **4.** Across all cells, the most common enriched pathways are specifically targeted to oxphos (in 20 different cell types, with increased expression of most subunits of mitochondrial complexes I-IV), DNA repair (8 cell types), Myc Targets V1 (7), the ROS pathway and EMT (4 each). The strong positive enrichments for oxphos suggests a major reconfiguration of the electron transport chain under the IH stimulus while downregulation of genes encoded by mitochondrial DNA in endothelial cells (LEC, AEC) and cells of the alveolocapillary unit (AF2, AT1, Cap1&2, AF1) suggest a specific mitochondrial stress. Finally, it is relevant to highlight that these responses occur while the mice were still being exposed to the hypoxic stimulus.

The cells showing the strongest response to IH in terms of DEGs are key elements that determine lung functions in health and disease. Although the purpose of the airway smooth muscle is ill-defined (34), many believe that its dysregulation in terms of enlargement (hyperplasia and cellular hypertrophy), extracellular matrix synthesis, and contractile capacity contribute to excessive airway narrowing in respiratory disorders such as asthma (35). While the number of ASMC have not increased substantially during IH exposures (it even slightly decreases, but our experimental approach precludes firm conclusion on the significance of this effect) the fact that there are specific enrichments for EMT and myogenesis (Hallmark) indicates responses that align with progression towards a phenotype associated with airway dysfunctions. Among the most important upregulated DEG that contribute to the EMT enrichment, several genes encoding collagen chains type 1 (col1a1, col1a2), 3 (col3a1), 5 and 6 (col5a2, col6a3, col6a2) and biglycan are components of the extracellular matrix, typically hypertrophied in asthmatic airways (36) - (cf supplemental data 11). Also noteworthy, is the increased expression of the transforming growth factor receptor Tgfb3 that plays a notable role in collagen synthesis during wound healing and pathological processes leading to fibrosis (37) and contributes to the selective enrichment of the EMT in ASMC.

Another striking feature of the response to IH exposure is the drastic reduction in the number of lymphatic endothelial cells (from 277 in Nx mice to 66 cells in IH), concomitant with fewer immune cells (B cells, cDC2, CD8, NK, CD4, neutrophils and iMOM). While our experimental design requires caution when comparing cell proportions between IH and Nx, this pattern – that is apparently coordinated among several cell types rather than being random changes – suggests a specific response shared among immune and LEC cells. Indeed, the lymphatic system in the lungs is a key component of the adaptive immune response, allowing trafficking of immune cells (dendritic cells and leukocytes) to the lymph nodes where the immune response to infection and inflammation is coordinated (38). Lymphatic drainage of accumulated fluids is also a key element of healthy lung homoeostasis, a striking example of this function being the decreased lymphatic vessels density found in lung sampled after fatal asthmatic crises, presumably contributing to the establishment of severe airway edema and obstruction (39). Similarly, lymphatic functions appear to be dysregulated in COPD (increased lymphatic density around alveolar spaces - (40)) and lung fibrosis (lymphatic density positively associated with fibrotic collagen and disease severity - (41)). Although the precise mechanisms underlying these associations remain unclear (38), it is noteworthy that pulmonary fibrosis is associated with severe histologic damage to lymphatic vessels presumably leading to drastic lymphatic dysfunctions (42). In mice, specific deletion of pulmonary lymphatic vessels results in chronic inflammation, alveolar enlargement, and profound hypoxemia with arterial oxygen saturation slightly above 80%, which are clear features of emphysema (43). Correlating with the drastic reduction of individuals LEC, IH exposure induced a profound regulation of the gene expression pattern in LEC, with significant enrichments for the EMT, DNA repair, TGFβ signaling, and oxphos. While ASMC and LEC share a similar enrichment for EMT, the genes contributing to this enrichment are different between the two cell types. LEC show specific upregulation of TGFβ1, and a host of genes encoding proteins that are associated with the extracellular matrix such as dystonin (Dst), Peptidyl-prolyl cis-trans isomerase B (Ppib), laminin subunit gamma-1 (Lamc1), fibulin 5 (Fbln5), col4a1 and elastin (Eln), suggesting major reorganization at this level (see supplemental data 12). Enrichment of the TGFβ signaling pathway in LEC was related to specific upregulation of TGFβ1/Smad1/Smurf1 (see supplemental data 13), which are also key components of fibrotic responses in the lungs (44).

In direct contrast with the drastic reduction of individual LEC, IH exposure induces a small increase in the AEC count (from 34 cells in Nx to 58 cells in IH), but we cannot draw a firm conclusion from these data due to our experimental constraints. AEC contribute to the development of PAH and exhibit a strong vasoconstrictive response under hypoxic conditions. Additionally, intermittent hypoxia promotes the muscularization of small pulmonary arteries and induces right ventricular hypertrophy, both of which are indicative of elevated pulmonary arterial pressure (9). The predominant response in AEC (DEG count), is a downregulation of gene expression, but without specific negative enrichment using the Hallmark collection of gene sets. Extending the GSEA to the GO collection showed negative enrichments related to the Notch pathway, angiogenesis, and response to TGFβ. Among the genes contributing to these enrichments are Notch1, Notch4, Hey1 (downstream in the Notch signaling pathway), Tgfb3, and Xbp1 (see supplemental data 14). The Notch signaling pathway is a major contributor to lung development and is involved in most respiratory diseases, contributing to the development of PAH, COPD, and fibrosis (45). Mechanistically, Notch1 appears to be downregulated in endothelial cells in COPD patients, and Notch overexpression reduces apoptosis (46).

A closer look to the alveolar-capillary unit cells (AT1, AT2, CAP1, CAP2, AF1, AF2, and AM) reported an apparent trend of coordinated expansion of the alveolar surface. An expanded AT1 population, whose ultra-thin cytoplasm allows gas exchange, could reflect enlarged functional respiratory surface, which is consistent with previous data showing that IH exposure increases tidal volume, inspiratory capacity and alveolar surface area in mice (4, 9), while inducing cellular proliferation (9). Interestingly, we previously reported that larger lungs and expanded alveolar surface area are also observed in mice living at high altitude, associated with elevated arterial O_2_ saturation and adaptive physiological responses (47, 48). Altogether, these data suggest the existence of a broad, and striking, adaptative strategy of lung growth and expansion of the alveolar surface in response to decreased oxygen availability in mice. AT1 cells roughly double in number (219 Nx-cells vs 415 IH-cells), while AT2 cells increase more modestly (86 Nx-cells vs 103 IH-cells). In parallel, we see a relative decrease in quiescent fibroblasts (AF1) and an increase in activated-like fibroblasts (AF2), hinting at extracellular-matrix deposition (49). While modest matrix production can stabilize and rebuild injured alveolar septa, excessive fibroblast activation risks fibrotic stiffening and impaired diffusion. Together, these cell-type adjustments (epithelial proliferation and maturation, macrophage recruitment, and fibroblast activation) may represent a mixed compensatory-pathological program aiming at increasing oxygen uptake at the cost of inflammation and fibrosis. Although our pooled-sample design precludes definitive claims of significance at the population level, the concordant directionality of these shifts aligns with known IH-induced lung remodeling, fibrogenesis, and inflammation in mice (6, 8, 9, 50).

A striking feature of the IH response, observed across a large variety of cells in the alveolar unit, as well as in other cell types, is the evident mitochondrial remodeling with enhanced expression of nuclear-encoded genes contributing to oxidative phosphorylation and selective reduction of the expression of some mitochondrial DNA genes in ASMC, LEC and AEC, in cells from the alveolar-capillary unit, and in some immune cells. Most genes of the complexes I-IV of the electron transport chain were upregulated in response to IH, and this was not related to the apparent expansion or reduction of the cell numbers even in those showing the most drastic increase (ex AT1) or decrease (LEC) of cell number. Rather, this response could be an efficient way to buffer the oxidative stress associated with the IH episodes (3) by increasing the flux of electrons that can be used to reduce oxygen in the mitochondria. Such responses can be particularly relevant during the reoxygenation phases, when most oxidative damage are likely to occur (51). In LEC, AEC and cells from the alveolar-capillary unit (AF2, AT1, cap1, cap2, AF1), the increased expression of nuclear genes coding for proteins of the electron transport chain is accompanied by a reduced expression of mitochondrial encoded genes. Such mito-nuclear imbalance can be triggered by specific metabolic stressors and is related to the mitochondrial unfolded protein response (31), contributing to the complex mitochondrial signalling networks (32, 33, 52).

Finally, it is particularly relevant to highlight that the responses observed under our experimental conditions largely differ from a previous report using a similar approach (13). One key experimental difference is the timing of lung sampling: while our animals have been exposed to IH for 3 hours (between 6:00 and 9:00 am) at the time of euthanasia and sampling, animals used in this previous study were maintained under normoxia overnight (standard for sleep apnea model in nocturnal rodents), but without being re-exposed to IH before tissue sampling. While it is beyond the scope of the present analysis to precisely describe these differences, this highlights one of the pressing needs to better understand to burden imposed by IH exposure. Indeed, cell-specific transcriptomic responses under IH exposure most certainly cycle between states of immediate responses (the present data set), and periods of recovery, after several hours under normoxia (previous data set in (13)). For examples, while we report that ASMC, LEC and AEC are among the cells that show strong responses to IH, this was not the case in the previous study. Furthermore, the mitochondrial response that we observe across most cell types was not found previously. Accordingly, it is tempting to speculate that transcriptomic regulations under typical IH models cycle through different phases, that could also have clinical relevance, and time course experiments could be useful to delineate early vs late response genes.

We conclude that in the lungs, ASMC, LEC and AEC show the most drastic transcriptomic responses under exposure to IH, responding to hypoxia in ways that could favor the development of lung diseases including asthma, fibrosis, PAH, and COPD. Most lung cells respond to IH exposure by selectively increasing the expression of genes coding for protein mitochondrial complexes I-IV and TCA cycle, suggesting a strategy to buffer the oxidative stress associated with hypoxic/reoxygenation episodes and mito-nuclear imbalance. Consistent with previous findings under chronic or intermittent hypoxia, there was also an apparent expansion of the alveolar-capillary unit, which might be a strategy for coping with frequent bouts of hypoxia. These responses differ from a previous data set obtained during a period of recovery from intermittent hypoxia, highlighting the need of defining the dynamic profiles of transcriptomic responses to further our understanding of pathological responses in this model of sleep apnea.

## Supporting information

see supplemental data 6A

see supplemental data 12

see supplemental data 13

see supplemental data 14

## Acknowledgements

Study funded by CIHR (PJX 192140) and Fondation de l’Institut Universitaire de Cardiologie et Pneumologie de Québec.

