## supplemental table 1 for "Cell-Specific Transcriptomic and Mito-Nuclear Imbalance in Lungs Under Intermittent Hypoxia in Adult Male Mice"

### Supplemental Figures 1-4

### Supplemental Tables 1-3

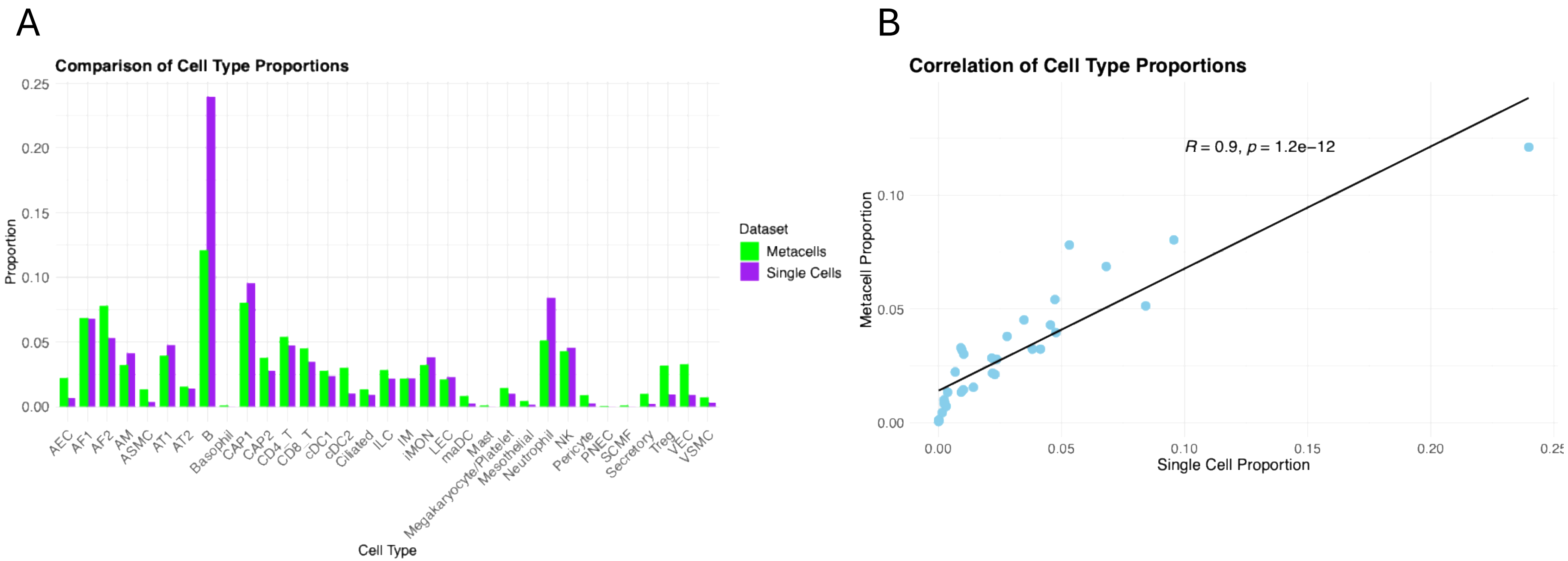


**Supplemental Figure 1: Comparison of cell type proportions between single-cell and aggregated supercells (metacells).**
A) Comparison of individual cell type proportions side by side for each dataset: single cell (purple) and metacells (green). B) Graphical representation of the correlation between single cell and metacell proportions. The correlation coefficient was calculated using Pearson's correlation.

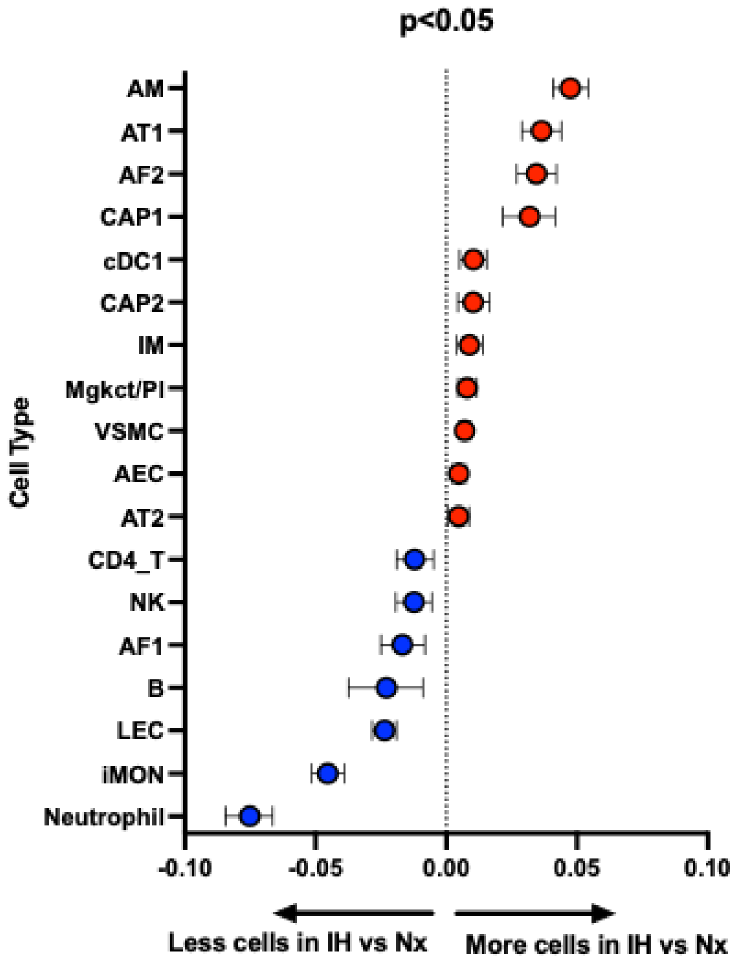


**Supplemental Figure 2: Bootstrap-estimated differences in cell type proportions between IH and Nx.** Only cell types with $p < 0.05$ (bootstrap) are shown. Note: due to pooling of biological replicates, results reflect cell-level variability and should be interpreted cautiously.

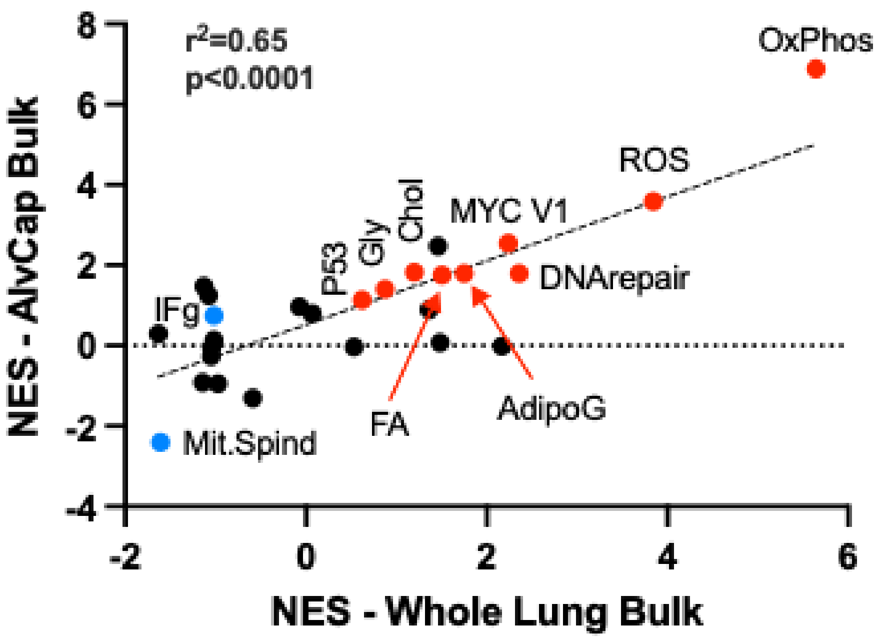


**Supplemental Figure 3: Correlation of significantly enriched gene sets (Hallmark collection, FDR < 0.1, either in the whole lung or in the AlvCap bulk) in response to IH between the pseudo bulks for the whole lung and the AlvCap unit.** Pathways that are common (FDR < 0.1) in both pseudo bulks are highlighted (red and blue). OxPhos: oxidative Phosphorylation. ROS: Reactive oxygen species pathway. MYCV1: MYC Targets V1. Chol: Cholestrol Homeostasis. Gly: Glycolysis. P53: P53 pathway. AdipoG: Adipogenesis. FA: Fatty acid metabolism. Ifg: Interferon gamma response. Mit.Spind: Mitotic Spindle.

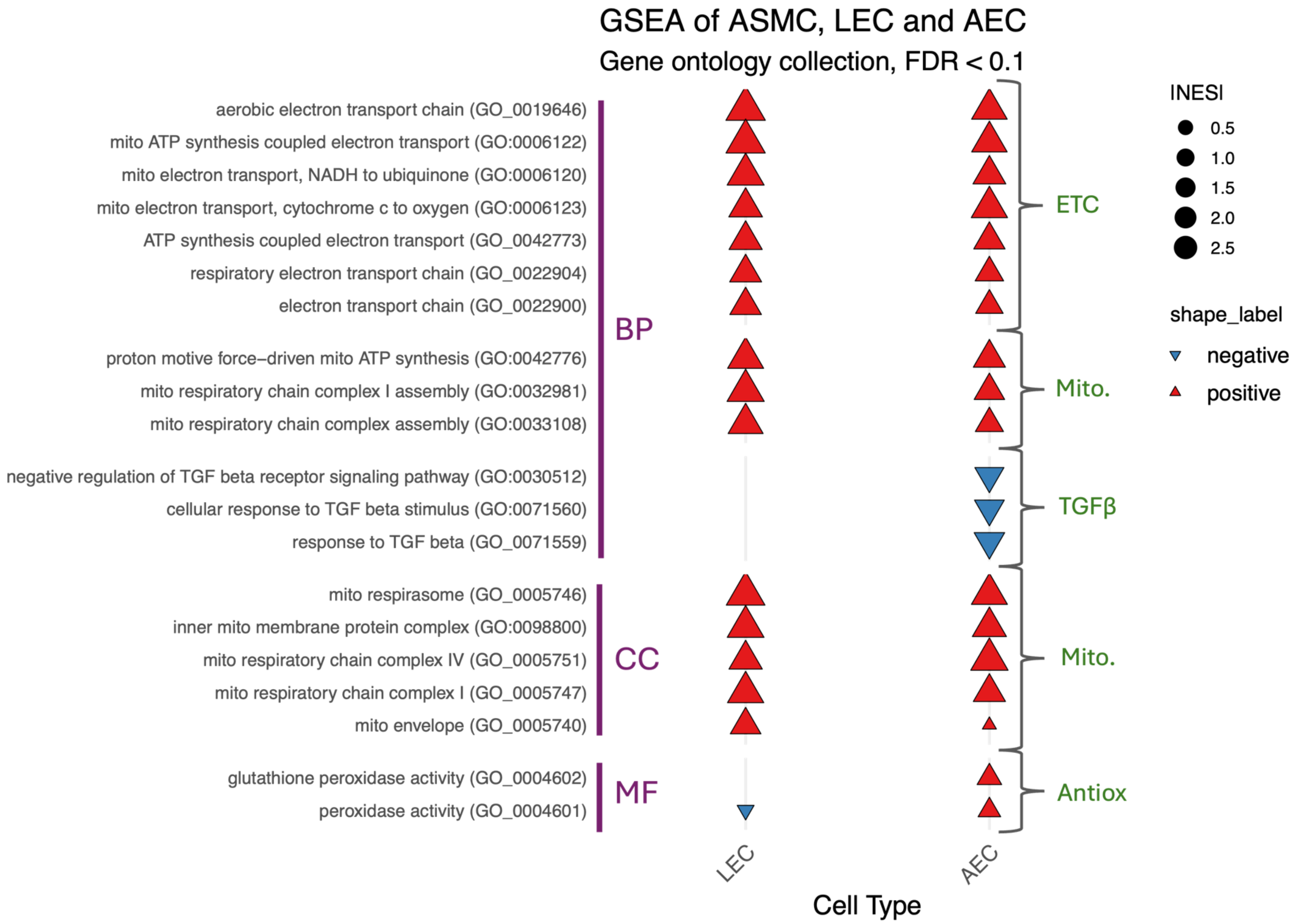


**Supplemental figure 4: Gene set enrichment analysis in ASMC, LEC and AEC (restricted to the mitochondrial, antioxidant and TGFβ genesets in the gene ontology collection).**
Gene sets were organized according to domains (BP: biological processes, CC: cellular components and MF: molecular functions) and restricted to pathways related to mitochondrial functions (mito.), electron transport chain (ETC), antioxidant activity (antiox.), and transforming growth factor beta (TGFβ) functions. Statistical threshold FDR<0.1.

Quality control filtering thresholds calculated by IsOutlier()

| Sample ID | nCount | | nFeature | | percent.mt | |
|---|---|---|---|---|---|---|
| | Min | Max | Min | Max | Min | Max |
| IH_mm10_vM23 | Not Applied | 22,970 | Not Applied | 7038 | Not Applied | 6.31% |
| NX_mm10_vM23 | Not Applied | 17,131 | Not Applied | 6181 | Not Applied | 4.93% |

**Supplemental table 1: Quality Control Filtering Thresholds for scRNA-seq Data**

Thresholds were derived using isOutlier() from the scater package, which identifies outliers based on robust statistical measures (median ± n × MAD). Negative lower thresholds were not applied as they are non-restrictive and biologically irrelevant for filtering.

| Group of cells | Proportion (%) | |
|---|---|---|
| | Nx | IH |
| **Immune cells** (B, AM, Neutrophil, CD4T, NK, CD8T, IM, cDC1, ILC, IMOM, megacaryocyte/platelet, Treg, cDC2, maDC, Basophil, Mast) | 66% | 55% |
| **alveolar epithelium-capillary interface cells** (CAP1, CAP2, AT1, AT2, AF1, AF2, AM) | 28% | 43% |
| **Total Immune cells + alveolar interface cells** | 94% | 98% |

**Supplemental Table 2:** Proportions of immune cell and cell composing the alveolo-capillary unit in the lungs in response to normoxia (Nx) or intermittent hypoxia (IH).

| CellType | Gene | Log2FC | q_value |
|---|---|---|---|
| ASMC | mt-Atp8 | -3.38 | 0.0003 |
| | mt-Co1 | -0.75 | 0.0033 |
| | mt-Co3 | -0.77 | 0.0068 |
| | mt-Nd2 | -0.86 | 0.0286 |
| | mt-Nd4 | -0.76 | 0.0101 |
| | mt-Nd4l | -2.57 | 0.0013 |
| LEC | mt-Atp6 | -0.81 | 0.0001 |
| | mt-Co1 | -1.08 | <0.0001 |
| | mt-Co2 | -0.99 | <0.0001 |
| | mt-Co3 | -0.96 | <0.0001 |
| | mt-Cytb | -0.77 | 0.0013 |
| | mt-Nd1 | -1.24 | <0.0001 |
| | mt-Nd2 | -0.69 | 0.0350 |
| | mt-Nd3 | -1.62 | 0.0305 |
| | mt-Nd4 | -0.71 | 0.0022 |
| | mt-Nd4l | -1.53 | 0.0358 |
| | mt-Nd5 | -1.42 | 0.0073 |
| AEC | mt-Atp8 | -2.00 | 0.0212 |
| | mt-Nd6 | -2.01 | 0.0159 |

| CellType | Gene | Log2FC | q_value |
|---|---|---|---|
| AF1 | mt-Atp8 | -1.35 | 0.0031 |
| | mt-Nd6 | -1.33 | 0.0028 |
| AF2 | mt-Atp8 | -1.27 | 0.0042 |
| | mt-Nd4l | -1.09 | 0.0031 |
| AT1 | mt-Atp8 | -1.94 | 0.0020 |
| | mt-Nd6 | -2.51 | <0.0001 |
| CAP1 | mt-Atp8 | -1.93 | <0.0001 |
| CAP2 | mt-Atp8 | -2.37 | 0.0001 |

| | | | |
|---|---|---|---|
| B | mt-Atp8 | -1.46 | <0.0001 |
| | mt-Nd4l | -0.90 | 0.0025 |
| | mt-Nd6 | -1.34 | <0.0001 |
| CD4 T | mt-Atp8 | -1.64 | 0.0029 |
| | mt-Nd4l | -1.33 | 0.0041 |
| | mt-Nd6 | -1.97 | 0.0001 |
| CD8 T | mt-Nd6 | -1.76 | 0.0069 |
| Treg | mt-Atp8 | -2.15 | 0.0065 |
| VEC | mt-Atp8 | -1.76 | 0.0288 |
| cDC1 | mt-Atp8 | -1.93 | 0.0276 |
| | mt-Nd4l | -1.51 | 0.0484 |
| iMON | mt-Nd4l | -1.52 | 0.0489 |
| | mt-Nd6 | -1.97 | 0.0174 |

**Supplemental Table 3:** Significant changes in mtDNA-encoded gene expression (|log2FC| > 0.5 & q < 0.05).

Supplemental table 3
